# QUALIS: The journal ranking system undermining the impact of Brazilian science

**DOI:** 10.1101/2020.07.05.188425

**Authors:** Rodolfo Jaffé

## Abstract

A system called QUALIS was implemented in Brazil in 2009, intended to rank graduate programs from different subject areas and promote selected national journals. Since this system uses a complicated suit of criteria (differing among subject areas) to group journals into discrete categories, it could potentially create incentives to publish in low-impact journals ranked highly by QUALIS. Here I assess the influence of the QUALIS journal ranking system on the global impact of Brazilian science. Brazil shows a steeper decrease in the number of citations per document since the implementation of this QUALIS system, compared to the top Latin American countries publishing more scientific articles. All subject areas showed some degree of bias, with social sciences being usually more biased than natural sciences. Lastly, the decrease in the number of citations over time proved steeper in a more biased subject area, suggesting a faster shift towards low-impact journals. Overall, the findings documented here suggest that the QUALIS system has undermined the global impact of Brazilian science, and reinforce a recent recommendation from an official committee evaluating graduate programs to eliminate QUALIS. A system based on impact metrics could avoid introducing distorted incentives, and thereby boost the global impact of Brazilian science.

## Introduction

In 1998 the Brazilian agency responsible for establishing criteria for evaluating the performance of higher education institutions (CAPES) launched a journal ranking system called “QUALIS”, which classified journals according to their distribution (local, national or international) and their quality within subject areas (A, B and C) (Andrade & Galembeck 2009). In 2009 this system was replaced by a new QUALIS (currently in use), which uses a complicated suit of criteria (differing among subject areas) to group journals into eight discrete categories (A1, A2, B1, B2, B3, B4, B5 and C) (Andrade & Galembeck 2009,Andriolo et al. 2010). The same journal can thus be ranked differently by different subject areas, and ranking criteria include different impact factor metrics, the proportion of journals in each category, the relevance or prestige of journals within subject areas, the number of issues published per year, the publishers, the need to support certain Brazilian journals, among others (a full explanation of the criteria employed by each subject area is available in Portuguese at: https://sucupira.capes.gov.br/sucupira/public/consultas/coleta/veiculoPublicacaoQualis/listaConsultaGeralPeriodicos.jsf; see also Appendix 1). QUALIS rankings are updated every four years by a group of academics from each subject area, and are used to evaluate the scientific production of graduate programs from higher education institutions in the following quadrennial (the last ranking was made with data from 2013-2016 and is being used to evaluate scientific production between 2017-2020). The system has a strong impact on Brazilian science, given that the distribution of funding resources and departmental fellowships are conditioned on the number of papers published in the highest QUALIS categories.

Although the QUALIS system has been subject to substantial criticism (da Silva 2009, Andriolo et al. 2010, Fernandes & Manchini 2019, Ferreira et al. 2013, Rocha-e-Silva 2009b), no systematic cross-subject area assessment has been yet performed to quantify its influence on the global impact of Brazilian science. This is surprising considering the system could create incentives to publish in low-impact journals ranked highly by QUALIS, thereby resulting in a decreased global impact. A steeper decrease in the number of citations per article (a measure of impact) since the implementation of the new QUALIS system, would indicate that QUALIS has actually undermined the impact of Brazilian science. However, because QUALIS criteria to rank journals differ between subject areas, some areas are expected to be more biased than others. We could thus anticipate that the relative decrease in the number of citations per article would be affected by the level of bias. Here I test these predictions.

## Materials and Methods

My aim was to assess the influence of the QUALIS journal ranking system on the global impact of Brazilian science. To this end I tested three specific predictions:

### 1) There has been a steeper relative decrease in the overall number of citations per document since the implementation of the new QUALIS system in 2009

Article citations are expected to increase with with article age, as older articles accumulate more citations than younger ones. When we plot citations over time, we expect to see a decrease in the number of citations, because more recent articles receive less citations tan older ones. However, this decline should be much slower (more gradual) for high-impact articles, since they usually accumulate citations faster and keep accumulating them for a longer time period. On the other hand, low-impact articles are expected to show a steep decline in the number of citations over time, as they need more time to accumulate citations and usually reach a plateau quickly, after which they stop accumulating citations. The shape of these plots, thus allows comparing the relative impact of articles, subject areas or countries. Here I plotted the total number of citations per documents (combining all subject areas) against time, using data from Scimago’s yearly country rankings between 2000 and 2019 (https://www.scimagojr.com/countryrank.php?year=2019&region=Latin%20America). The number of citations per document represents average citations to documents published during each year, so it is not biased by the absolute number of publications. I chose the top five Latin American countries publishing more scientific journal articles to perform this comparison (according to Scimago’s 2019 country rankings: Argentina, Brazil, Chile, Colombia, and Mexico). To assess these effects numerically I also calculated the Area Under the Curve (AUC) for each country, between 2000 and 2009 (pre QUALIS) and between 2010 and 2019 (post QUALIS). A steeper relative decrease in the number of citations since 2009 (resulting in a lower AUC) would indicate a negative effect of QUALIS in the global impact of Brazilian articles (i.e. articles being less cited).

### 2) Since QUALIS criteria to rank journals differ between subject areas, some areas are more biased than others

I used the proportion of journals indexed in the Scopus database in each of the QUALIS subject areas as a first proxy of bias. I chose the List of Scopus Index Journals (36,500 journals) because it contained more journals than Scimago Journal Rank (26,199 journals) or InCites Journal Citation Reports (12,300 journals). Scopus data for February 2019 was downloaded from this site: https://www.researchgate.net/publication/330967992_List_of_Scopus_Index_Journals_February_2019_New (doi: 10.13140/RG.2.2.23454.79688). To be indexed by Scopus, journals should meet all of the following minimum criteria: a) Consist of peer-reviewed content and have a publicly available description of the peer review process; b) Be published on a regular basis and have an International Standard Serial Number (ISSN) as registered with the ISSN International Centre; c) Have content that is relevant for and readable by an international audience; and d) Have a publicly available publication ethics and publication malpractice statement (see a more detailed description of Scopus’s evaluation criteria here: https://www.elsevier.com/solutions/scopus/how-scopus-works/content/content-policy-and-selection).

I then employed Scopus’s CiteScore as a proxy of the journal’s realized global impact. Scopus’s CiteScore 2017 represents the number of citations received in 2017 to documents published in 2014, 2015 and 2016, divided by the number of documents published in 2014, 2015, and 2016. Since it employs a 3-year citation window, rather than the 2-year window of the traditional Impact Factor, it approaches better the QUALIS quadrennial classification. The last QUALIS ranking was made with data for 2013-2016, so I collected CiteScore 2017 for journals comprised in all QUALIS subject areas (49 subject areas, 27,619 journals, raw data is available in the CAPES website: https://sucupira.capes.gov.br/sucupira/public/consultas/coleta/veiculoPublicacaoQualis/listaConsultaGeralPeriodicos.jsf, and here: https://github.com/rojaff/qualis). I used the journal’s ISSN number to match both databases (QUALIS and Scopus), reading all values as text to avoid loosing leading zeros. I then ran a Kruskal-Wallis test to compare the overall variation in CiteScore across QUALIS categories, and used chi-squared values as a measure of the strength of this variation (this test was chosen considering the non-normal distribution of CiteScore). I also assessed the number of cases when lower QUALIS categories had a higher median CiteScore than preceding higher QUALIS categories (example: median of B1 > median of A2). Finally, I calculated the number of journals classified as A1 having a CiteScore below the median CiteScore for each subject area.

I thus calculated four bias metrics:

i. Proportion of QUALIS journals indexed by Scopus: Since a higher proportion of journals indexed by Scopus implies that more journals pass Scopus’s minimum eligibility criteria, subject areas with a larger proportion of indexed journals are expected to be less biased.
ii. Kruskal-Wallis chi-squared: Since higher chi-squared values indicate stronger differences in CiteScore between QUALIS categories, subject areas with higher chi-squared values are expected to be less biased.
iii. Cases where lower QUALIS > higher QUALIS: Since larger values (more such cases), indicate that more journals in lower ranking categories have a higher CiteScore than those of the preceding, higher ranking category, subject areas with larger values of this indicator are expected to be more biased.
iv. A1 journals with CiteScore below the median CiteScore for each subject area: Since the A1 category is supposed to contain the area’s top ranking journals, higher values (more journals) indicate more journals have been miss-classified as A1, so subject areas with higher values of this indicator are expected to be more biased.

### 3) The relative decrease in the number of citations per document is affected by the level of bias

I identified the top less and more biased subject areas according to the four bias metrics described above, using the lower (5%) and upper (95%) quantiles as cutoff values for each metric. I then chose two subject areas that were ranked in each of these top groups using more than one bias metric. The number of citations per document between 2009 and 2019 received by Brazilian journal articles belonging to these two subject areas were then plotted against time, using data for the most similar subject areas from Scimago (note that Scimago has its own subject area classification). To make the comparison extreme, I chose a more biased subject areas with a higher number of citations per documents in 2009 than those of the less biased subject area. All analyses were performed in R and data and scripts are publicly available in GitHub: https://github.com/rojaff/qualis.

## Results

While the number of scientific papers produced by Brazilian scientist has increased during the past two decades, since 2009 the number of citations per document has remained the lowest among the top five Latin American countries publishing more scientific papers (Fig. 1). Brazil showed a higher area under the curve (AUC) than Mexico before QUALIS was implemented (2000-2009), but the lowest among all the evaluated countries after QUALIS was implemented (2010-2019, Table 1). From all journals comprised in the QUALIS system (including all subject areas), 16,760 (61%) were not indexed by Scopus (Table S2 shows the full list of non-indexed journals comprised in QUALIS). Across all subject areas the proportion of journals indexed by Scopus ranged between 6% and 73% (median = 50%, Fig. 2). I was able to retrieve CiteScore for 9,985 of the journals comprised in QUALIS (36%), and only 3% of the QUALIS journals indexed by Scopus did not contain a CiteScore (Fig.2, see absolute values in Table S1). The distribution of journal’s CiteScore values across QUALIS categories showed a very large variation across subject areas (Fig. 3). Remarkably, all subject areas showed some degree of bias in at least one bias indicator (Tables 2 and S1, Figs. S1-S4). In general, subject areas belonging to the social sciences were among the top more biased, whereas those belonging to the natural sciences were among the top less biased, with a few exceptions (Table 2).

**Table 1:** Area Under the Curve (AUC) for the number of citations per document against time (year) in the five Latin American countries publishing more scientific journal articles. AUC was calculated before the implementation of QUALIS (years 2000-2009) and after (years 2010-2019).

| Country | AUC |  |
| --- | --- | --- |
|  | 2000-2009 | 2010-2019 |
| Chile | 247.11 | 109.56 |
| Argentina | 222.72 | 98.7 |
| Colombia | 205.34 | 83.89 |
| Mexico | 185.98 | 82.35 |
| Brazil | 192.08 | 77.89 |

**Table 2:** Top less and more biased subject areas according to four bias metrics (see methods for details). Each group is composed of the lower (5% quantile) or upper (95% quantile) subject areas. Subject areas in bold were ranked in these quantiles using more than one bias metric. Original QUALIS subject area names are shown (as written in their respective classification sheets) but their English translation can be found in Table S1.

| Bias metric | Top less biased | Top more biased |
| --- | --- | --- |
| Proportion of QUALIS journals indexed by Scopus | **astronomia_fisica**,<br>ciencias_biologicas_iii,<br>medicina_iii | **ciencias_da_religiao_e_teologia, direito, servico_social** |
| Kruskal-Wallis chi-squared | Interdisciplinar, medicina_i,<br>**medicina_ii** | **antropologia_arqueologia, ciencias_da_religiao_e_teologia, filosofia** |
| Cases where lower QUALIS > higher QUALIS | **astronomia_fisica**,<br>engenharias_iv, **medicina_ii**,<br>psicologia * | **antropologia_arqueologia, direito, educacao, servico_social** |
| A1 journals with CiteScore below the area median | **astronomia_fisica**, materiais,<br>servico_social | economia, enfermagem, ensino |

\* In this case I used the lower 10% quantile as cutoff since the 5% quantile did not contain any subject areas.

**Figure 1:**
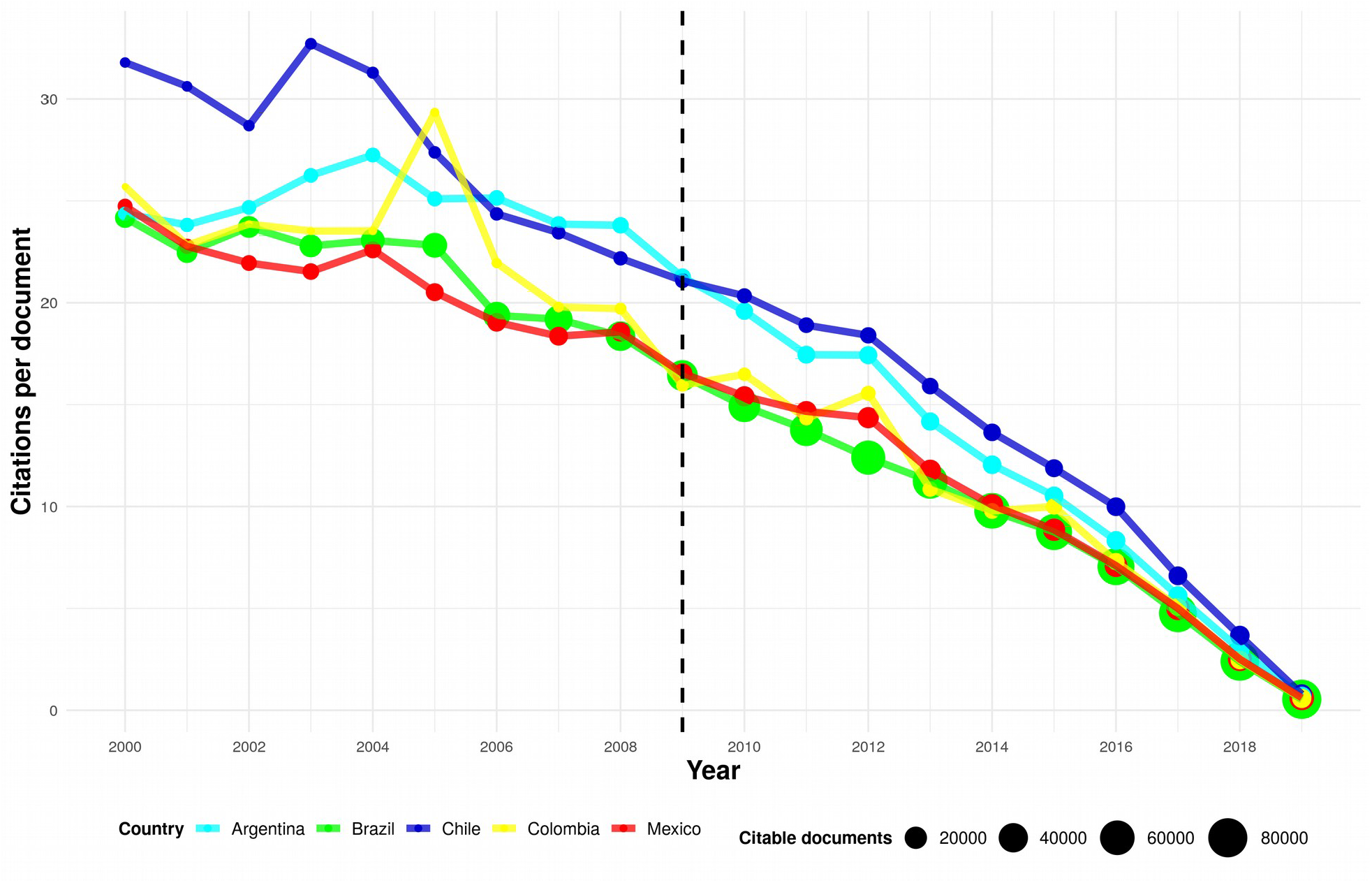
Number of citations per document between 2000 and 2019 in the five Latin American countries publishing more scientific journal articles (according to Scimago’s 2019 country ranking). Countries are identified by colors, while dot size represent the total number of citable documents. The vertical dashed line indicates the year when the new QUALIS system was introduced in Brazil.

**Figure 2:**
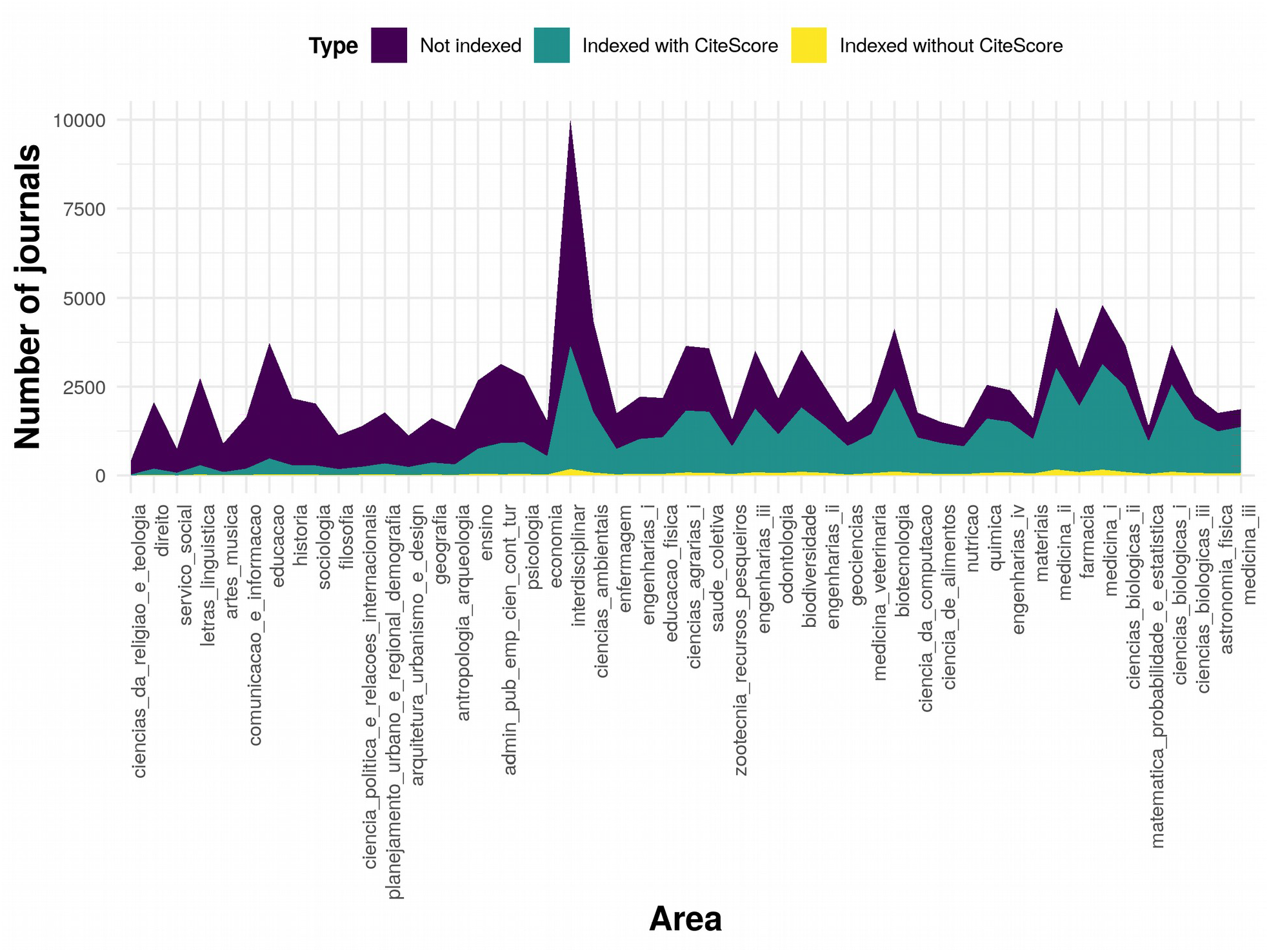
Number of journals not indexed by Scopus, indexed with available CiteScore 2017, and indexed without CiteScore 2017, across all 49 QUALIS subject areas. Original QUALIS subject area names are shown (as written in their respective classification sheets) but their English translation can be found in Table S1. Subject areas are sorted by the proportion of QUALIS journals indexed by Scopus.

**Figure 3a:**
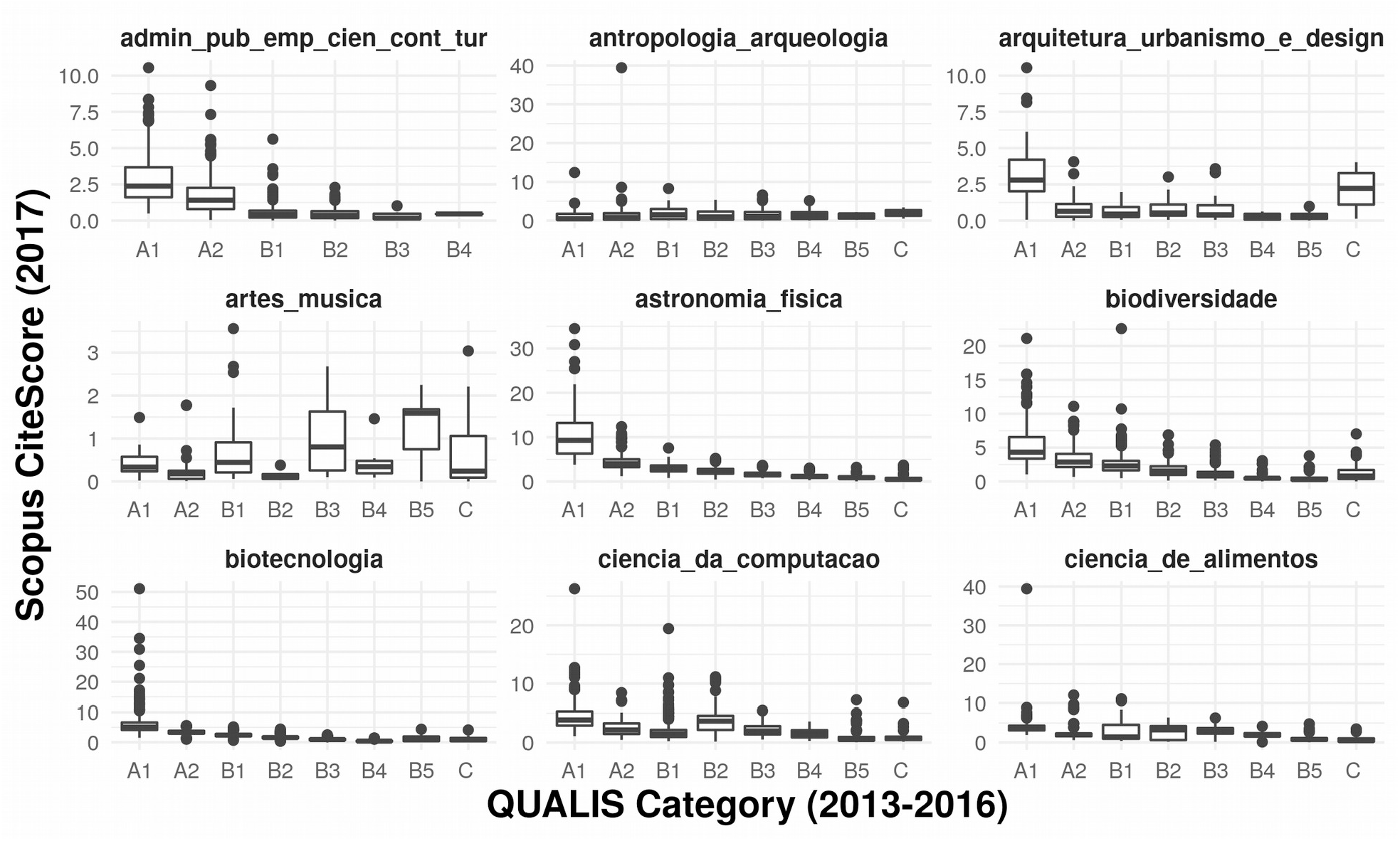
Scopus CiteScore variation across QUALIS categories in nine subject area. Original QUALIS subject area names are shown (as written in their respective classification sheets) but their English translation can be found in Table S1.

**Figure 3b:**
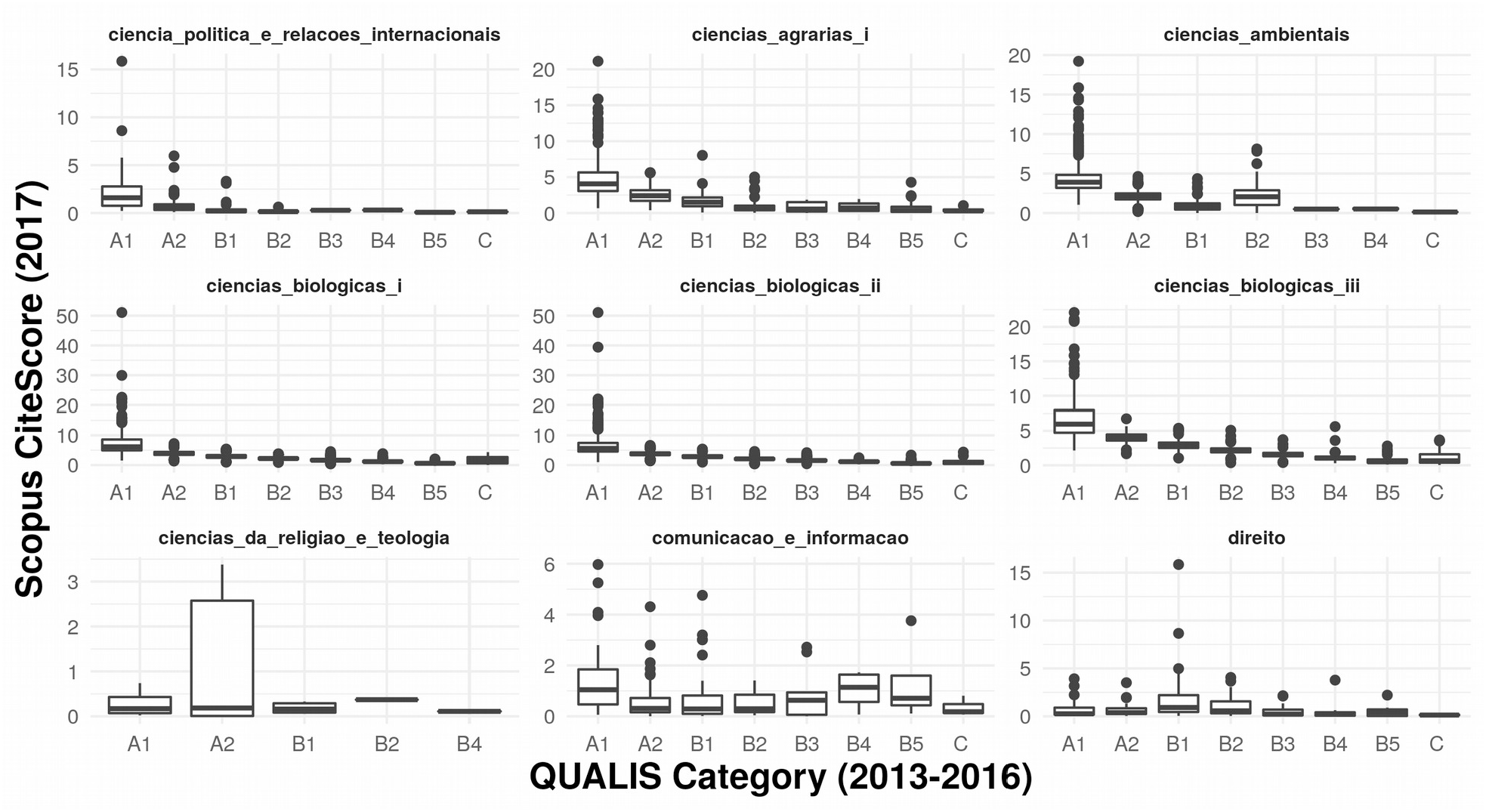
Scopus CiteScore variation across QUALIS categories in nine subject area. Original QUALIS subject area names are shown (as written in their respective classification sheets) but their English translation can be found in Table S1.

**Figure 3c:**
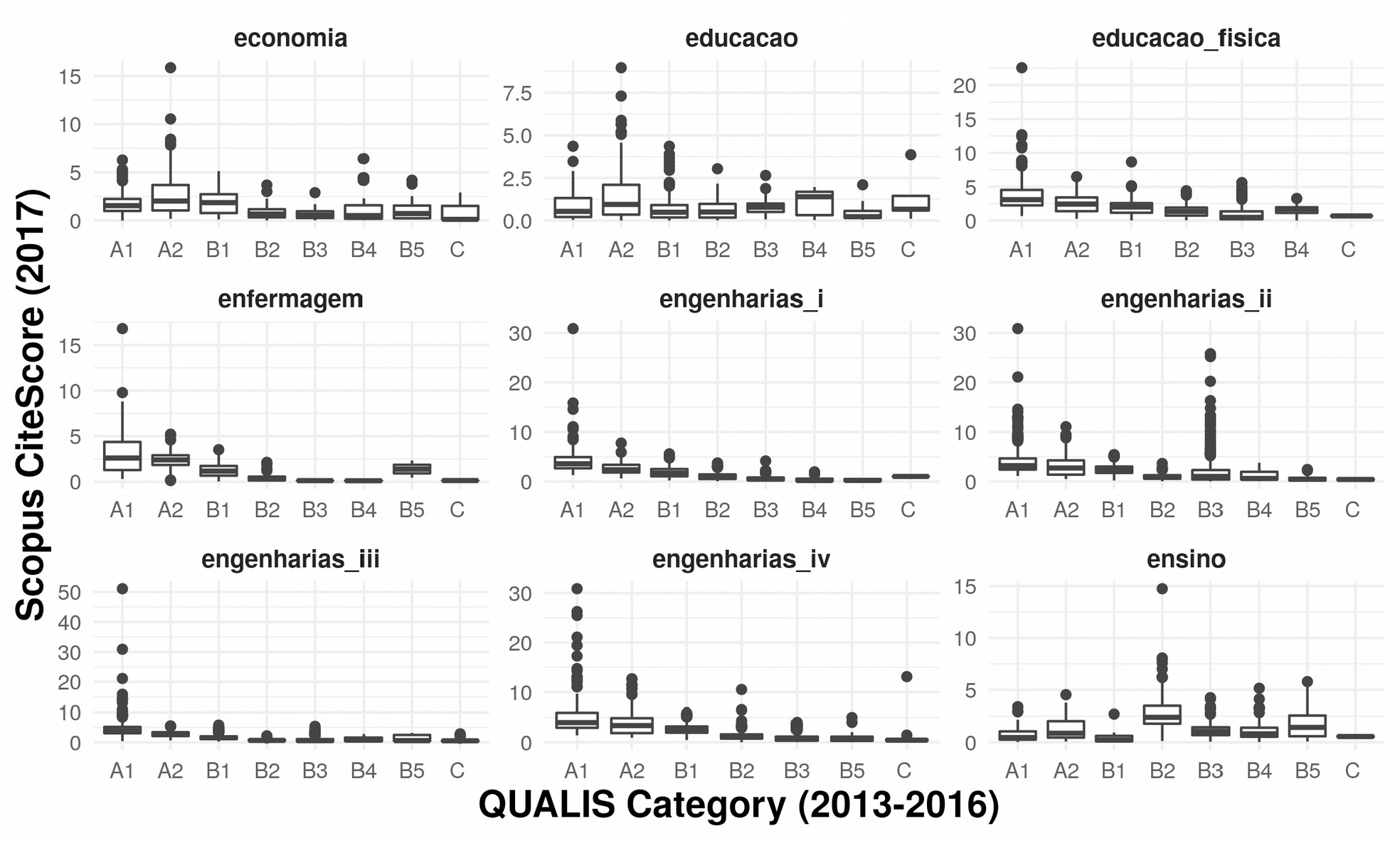
Scopus CiteScore variation across QUALIS categories in nine subject area. Original QUALIS subject area names are shown (as written in their respective classification sheets) but their English translation can be found in Table S1.

**Figure 3d:**
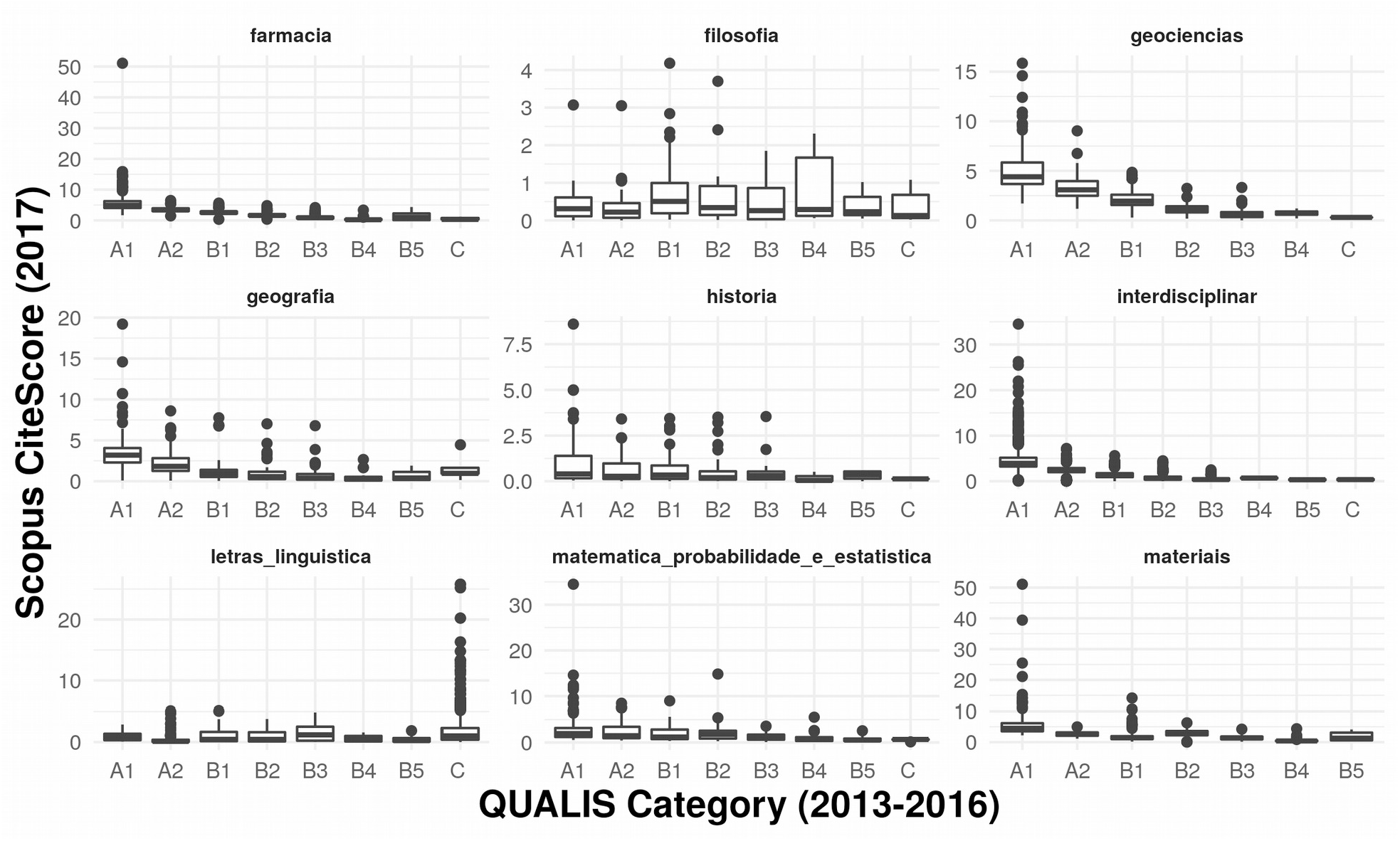
Scopus CiteScore variation across QUALIS categories in nine subject area. Original QUALIS subject area names are shown (as written in their respective classification sheets) but their English translation can be found in Table S1.

**Figure 3e:**
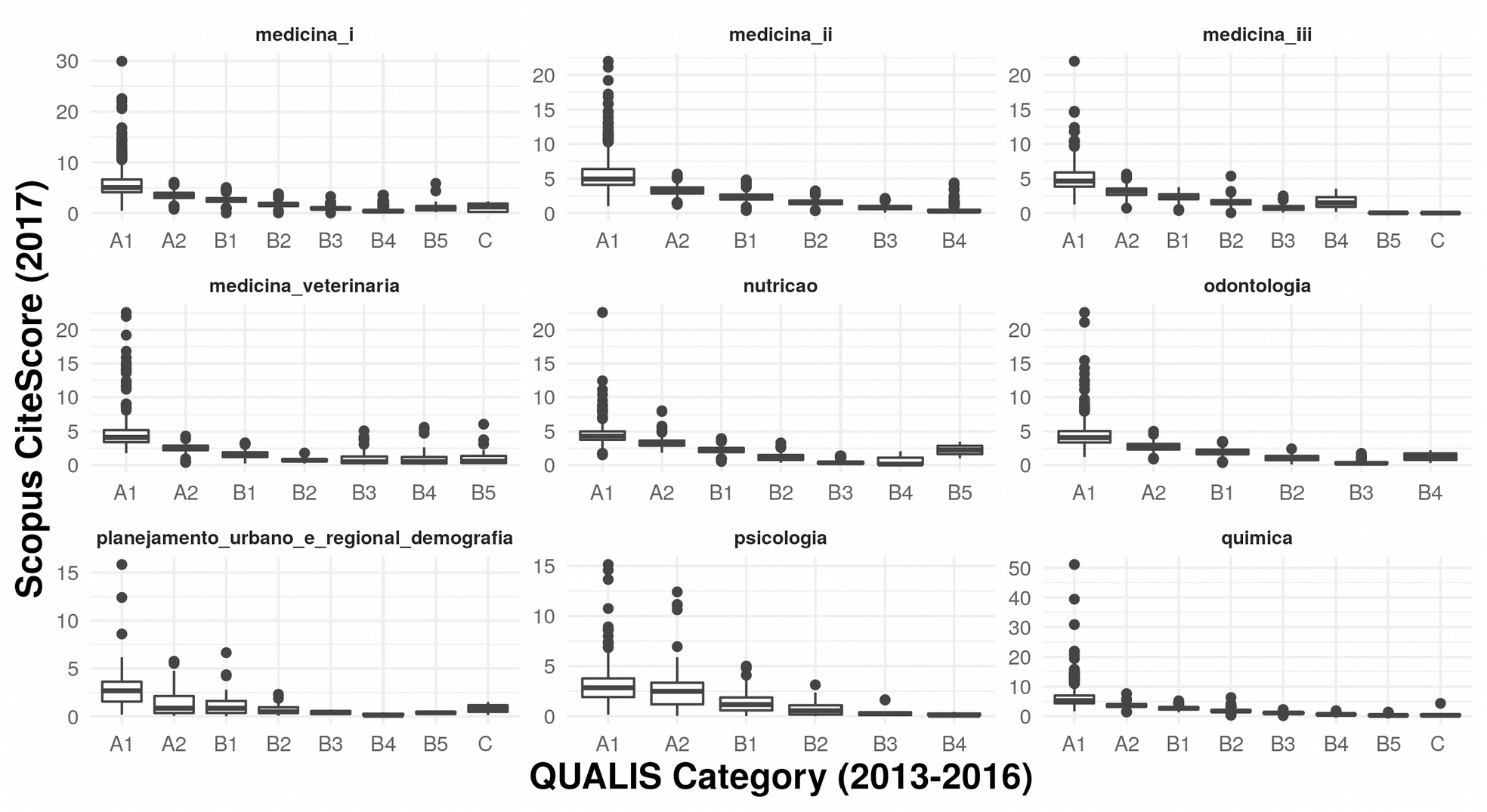
Scopus CiteScore variation across QUALIS categories in nine subject area. Original QUALIS subject area names are shown (as written in their respective classification sheets) but their English translation can be found in Table S1.

**Figure 3f:**
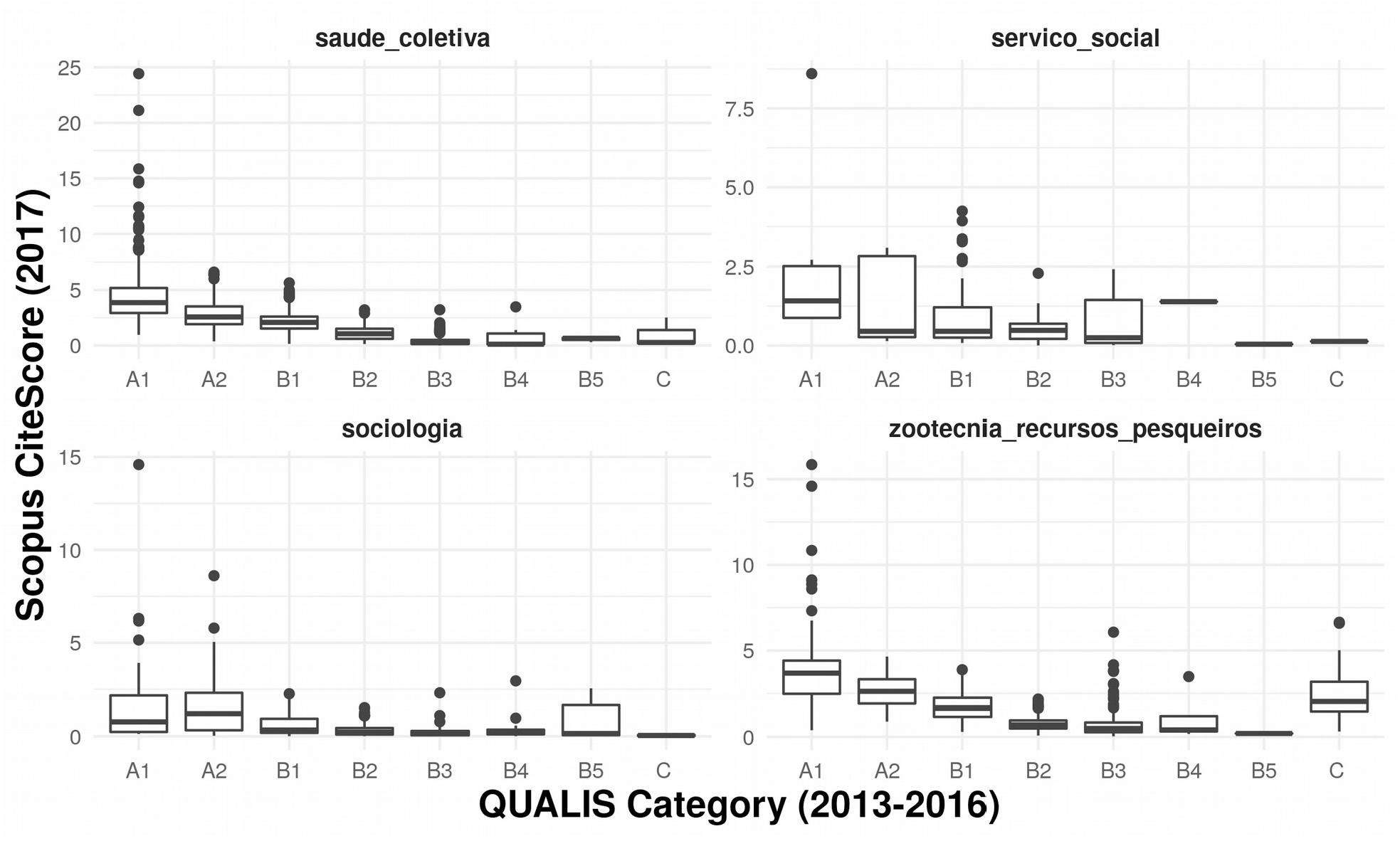
Scopus CiteScore variation across QUALIS categories in four subject area. Original QUALIS subject area names are shown (as written in their respective classification sheets) but their English translation can be found in Table S1.

The two selected subject areas that were ranked in the top less and more biased groups using more than one bias metric were “Astronomy and Physics” and “Social Work”, respectively. The number of citations per document received by Brazilian journal articles belonging to these two subject areas showed a progressive decrease in time, as expected. However, this decrease was much steeper in “Social Work” than in “Physics and Astronomy”, and even though “Social Work” had a higher number of citations per document in 2009, its area under the curve (AUC = 110.02) was smaller than that of “Physics and Astronomy” (AUC = 116.32; Fig. 4).

**Figure 4:**
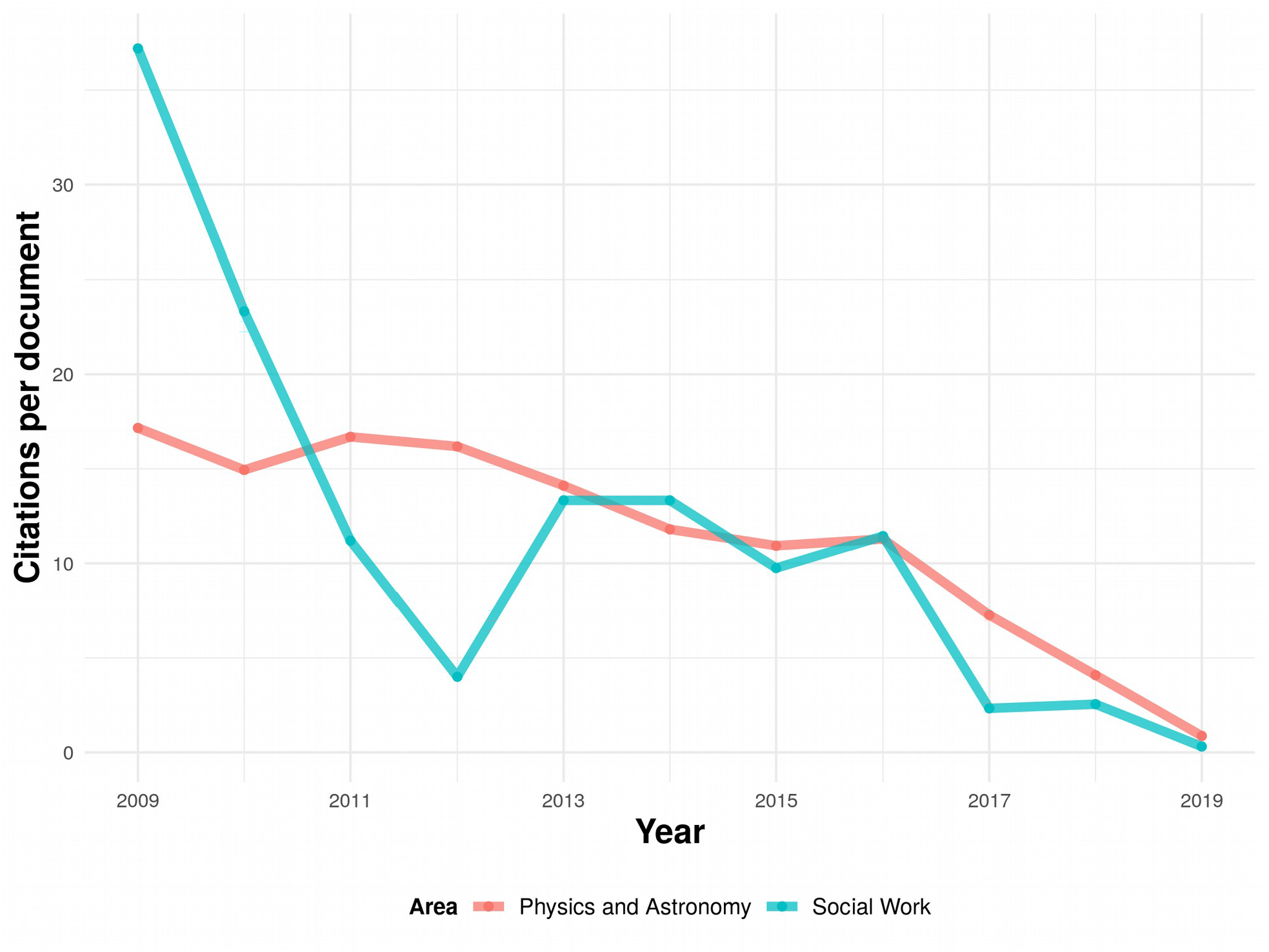
Number of citations per document received between 2009 and 2019 by Brazilian journal articles belonging to Scimago’s subject areas “Social Work” (AUC = 110.02) and “Physics and Astronomy” (AUC = 116.32).

## Discussion

Results reveal that the QUALIS system, originally intended to rank graduate programs from different subject areas and promote selected national journals, has been unable to increase the relative impact of Brazilian science, compared to other Latin-American countries, since its implementation in 2009. Moreover, all the subject areas making up the QUALIS system showed some degree of bias, with social sciences being usually more biased than natural sciences. Finally, the decrease in the number of citations over time was steeper in “Social Work” (a more biased subject area) than in “Physics and Astronomy” (a less biased subject area).

The steeper decline in the number of citations per document since 2009, compared to the top Latin American countries publishing more scientific papers, suggest that the QUALIS journal ranking system has created incentives to publish in low-impact journals ranked highly by QUALIS. For instance, changes in the QUALIS journal rankings have affected submission rates in journals like *Anais da Academia Brasileira de Ciências*: Submissions from Biological Sciences plummeted after this subject area downgraded the journal from A2 to B2 in 2013 (Kellner 2017). As faculty and graduate students are evaluated based on the number of papers they publish in journals that are highly ranked by QUALIS, they are more likely to select journals in the A categories with lower impact factors and higher acceptance rates (Aarssen et al. 2008). Over time, this system appears to have shifted publications towards low-impact journals ranked highly by QUALIS, thus undermining the global impact of Brazilian science. In contrast, in countries where scientists are evaluated based on the impact factor of the journals where they publish, the number of citations per document accumulate more quickly (so the curve shows a slower decrease over time). This effect is exemplified by Mexico and Colombia, which matched Brazil in number of citations per document in 2009 (when the current QUALIS system was implemented), but show a less abrupt fall since (Fig. 1).

Most of the journals comprised in the QUALIS system (61%) were not indexed by Scopus, and halve of all subject areas had less than 50% of its journals indexed in Scopus (Fig. 2). These results suggest that a substantial number of journals comprised in the QUALIS system do not meet the minimum eligibility criteria of the largest source-neutral database (Scopus). This is alarming, and reveals a need to set higher journal quality standards across all subject areas.

Although some subject areas were found to be more biased than others by the QUALIS system, all showed some degree of bias in at least one bias indicator. This result is surprising, and indicates that even in hard, quantitative areas, QUALIS journal ranks do not reflect the journal’s realized impact. In environmental sciences (top right plot in Fig. 3b), for example, category B2 has a higher median CiteScore than category B1, and there are journals classified as B2 showing a CiteScore above the A1 and A2 medians. Similar patterns are observed in many other subject areas, revealing that the multiple criteria used to create QUALIS journal ranks result in a mismatch between the perceived and the realized journal’s impact. Biases nevertheless appear to be more pronounced in the social sciences, suggesting a marked disregard for impact metrics (Table 2, Figs. S1-S4). Remarkably, in five subject areas (antropologia_arqueologia, ciencias_da_religiao_e_teologia, filosofia, historia, servico_social) CiteScore values did not significantly differ between QUALIS categories (Fig. 3, Table S1), indicating that the QUALIS rankings do not consider the journal’s impact factor at all.

Two of the least and most biased subject areas (“physics and astronomy” and “social work”, respectively) showed striking different patterns of citations over time, with social work exhibiting a sharp decline after 2009 (Fig. 4). This result indicates a rapid shift towards low-impact journals ranked highly by QUALIS in social work since 2009, resulting in an overall decrease in impact. In contrast, in physics and astronomy the QUALIS journal ranking follows the journal’s realized impact (CiteScore) more closely, so incentives are in place to publish in high-impact journals (also ranked highly by QUALIS). Perhaps thanks to these publications in high-impact journals, the number of citations per document accumulate more quickly (see right to left increase in Fig. 4). These findings reinforce that the QUALIS system implemented in 2009 is likely a major driver of the steeper overall decline in the number of citations per document since 2009, compared to the top Latin American countries publishing more scientific papers.

Overall, the findings documented here suggest that the QUALIS system has undermined the global impact of Brazilian science its implementation in 2009. Likewise, they reveal that a journal ranking system based on the realized impact of journals would avoid introducing distorted incentives, and thereby boost the global impact of Brazilian science (da Silva 2009). It is also difficult to justify QUALIS as a mean to promote national journals in the age of open-access and preprints (Andriolo et al. 2010, Kellner 2017, Rocha-e-Silva 2009a), and less so if it is at the expense of the global impact of Brazilian science (Ferreira et al. 2013). QUALIS was once considered a temporary strategy (da Silva 2009), and a recent report by CAPES has recommended it should not be used in the future any more, being replaced with “internationally established and broadly recognized metrics” (COMISSÃO ESPECIAL DE ACOMPANHAMENTO DO PNPG 2020). In another official communication (Ofício Circular n° 31/2020-GAB/PR/CAPES: http://uploads.capes.gov.br/files/OF_CIRCULAR_31-2020-GAB-PR-CAPES.pdf), CAPES recently informed the adoption of a new Reference QUALIS system, whereby journals will be ranked by a single mother subject area based on impact metrics like CiteScore, JCR, and *h*-index. However, whereas the Natural Sciences will rank journals based on these impact metrics, Humanities, Education and Public Health will be able to define their own criteria. The results presented here suggest that the best formula to boost the global impact of Brazilian science is to avoid the introduction of distorted incentives, using internationally recognized impact metrics to rank journals and evaluate graduate programs across all subject areas.

## Acknowledgments

I thank Klaus Jaffé for feedback and discussions that helped shape this manuscript. Marcelo Lattarulo Campos and two other anonymous referees helped improve earlier versions of this manuscript. I also thank Conselho Nacional de Desenvolvimento Científico e Tecnológico (CNPq) for a research productivity grant (301616/2017-5).

## Data Deposition

- All data used in this manuscript is publicly available and sources have been cited in the text.
- R scripts and data are publicly available in GitHub: https://github.com/rojaff/qualis
- Original QUALIS data was downloaded fromhttps://sucupira.capes.gov.br/sucupira/public/consultas/coleta/veiculoPublicacaoQualis/listaConsultaGeralPeriodicos.jsf
- Original Scimago data was downloaded from https://www.scimagojr.com/countryrank.php
- Original Scopus data was downloaded from https://www.researchgate.net/publication/330967992_List_of_Scopus_Index_Journals_February_2019_New

