## Supplemental Information for "QUALIS: The journal ranking system undermining the impact of Brazilian science"

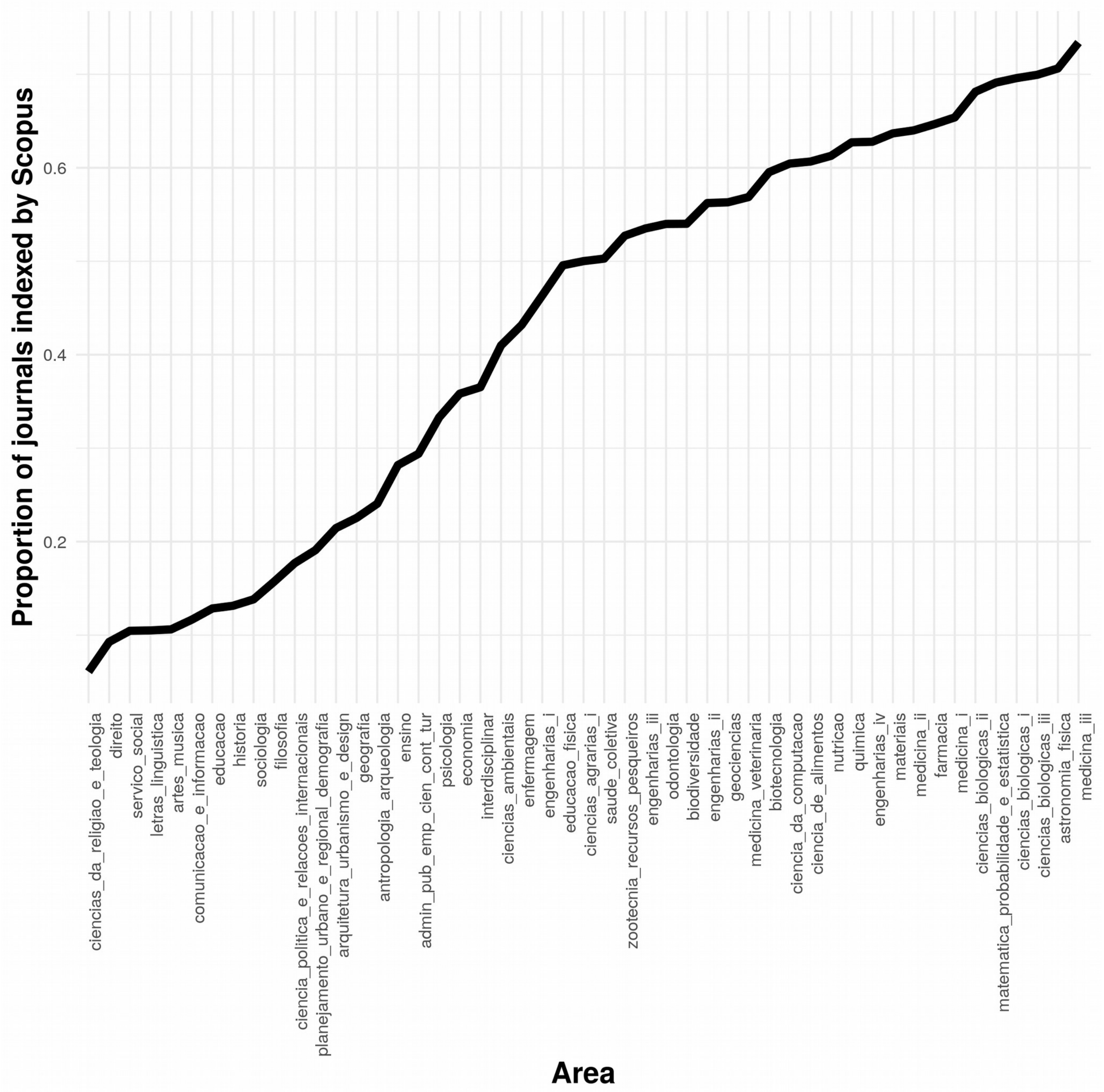


**Figure S1:** Proportion of journals indexed by Scopus across all 49 QUALIS subject areas. Original QUALIS subject area names are shown (as written in their respective classification sheets) but their English translation can be found in Table S1.

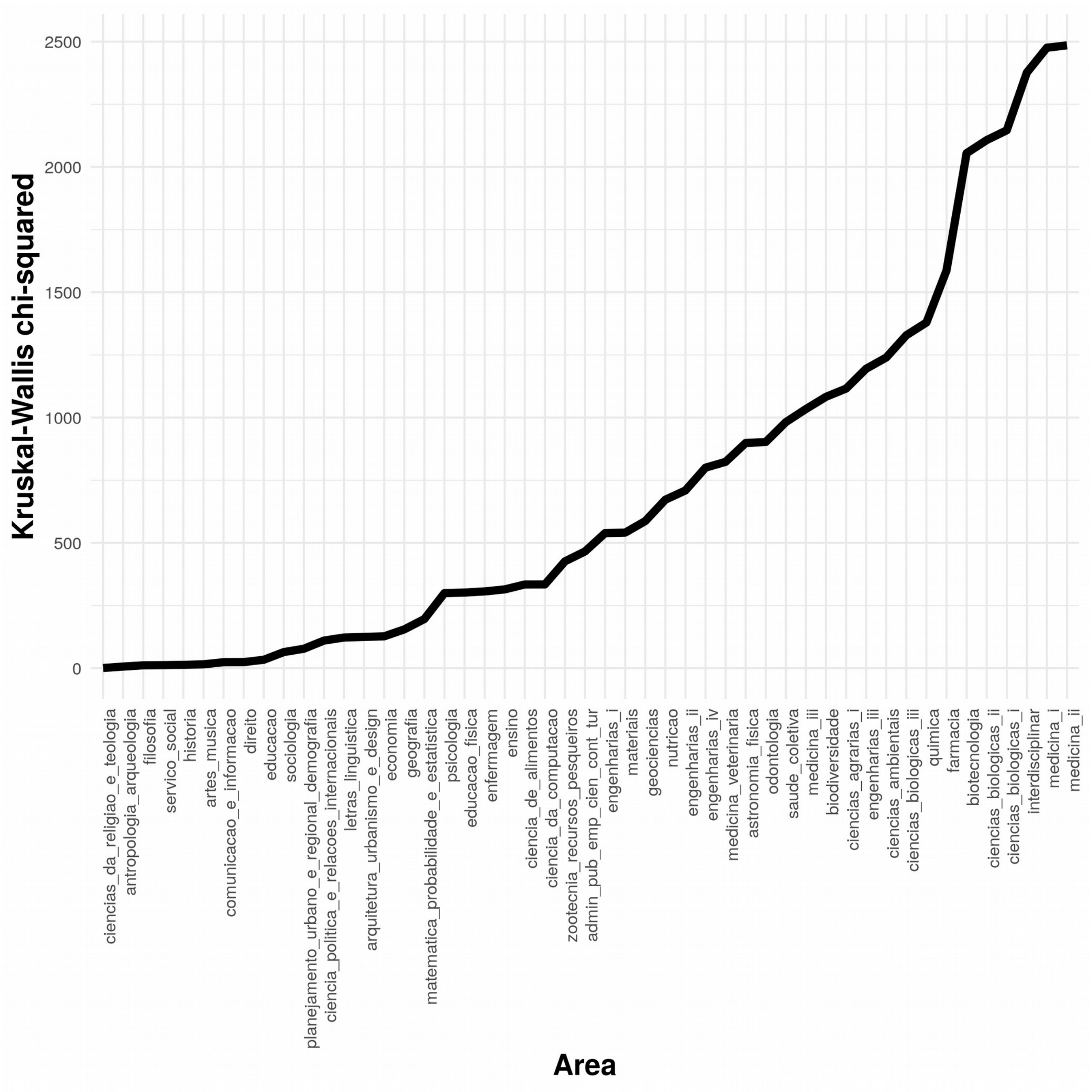


**Figure S2:** Kruskal-Wallis test chi-squared values across all 49 QUALIS subject areas. Tests assessed the variation of Scopus CiteScore across QUALIS categories in each subject area. Test statistics where significant ($p<0.05$) for all subject areas except the first four on the left side of the plot (see values in Table S1). Original QUALIS subject area names are shown (as written in their respective classification sheets) but their English translation can be found in Table S1.

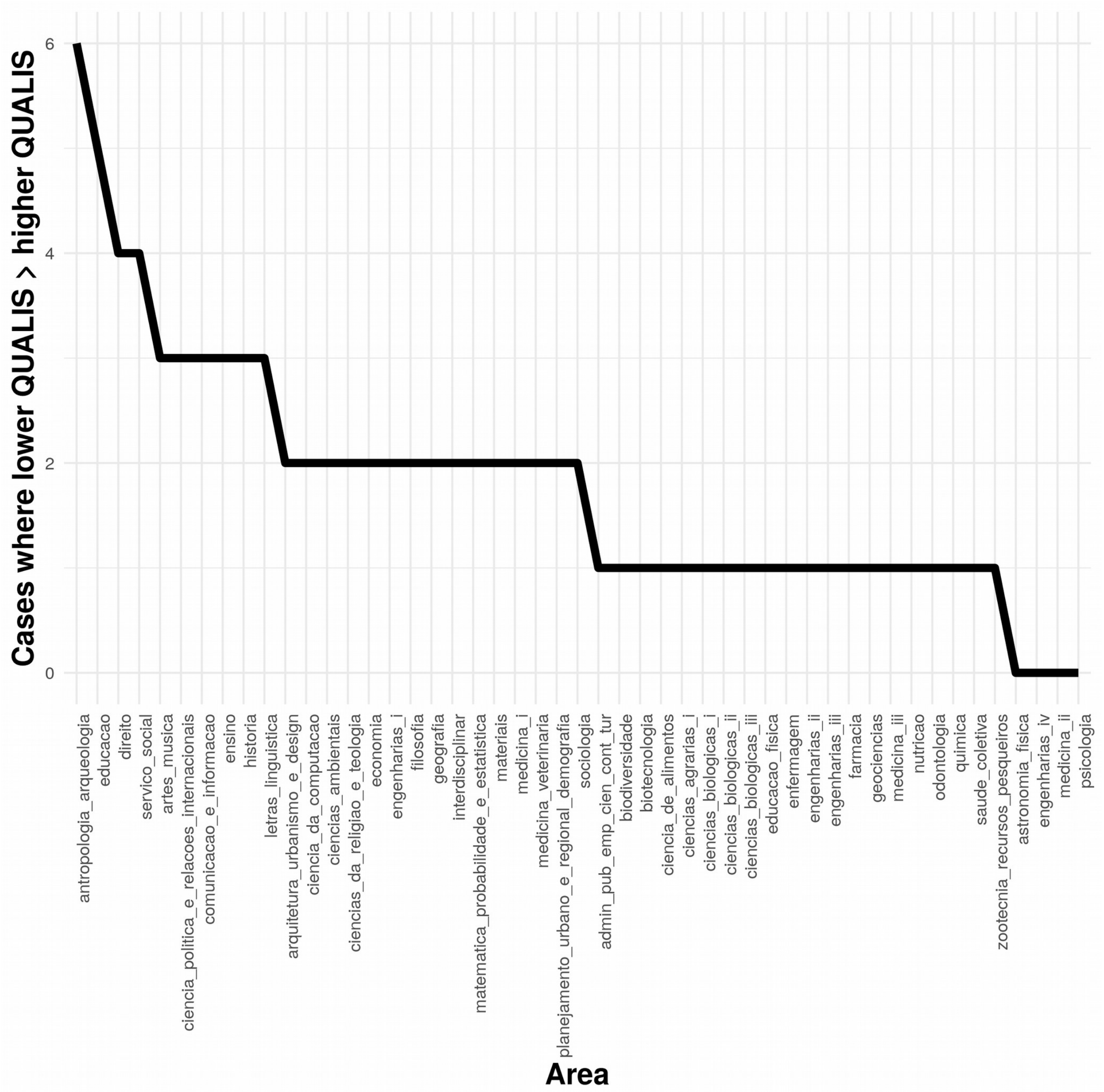


**Figure S3:** Number of cases where lower QUALIS categories had a higher median value than preceding higher QUALIS categories (example: median of B1 > median of A2) across all 49 QUALIS subject areas. Original QUALIS subject area names are shown (as written in their respective classification sheets) but their English translation can be found in Table S1.

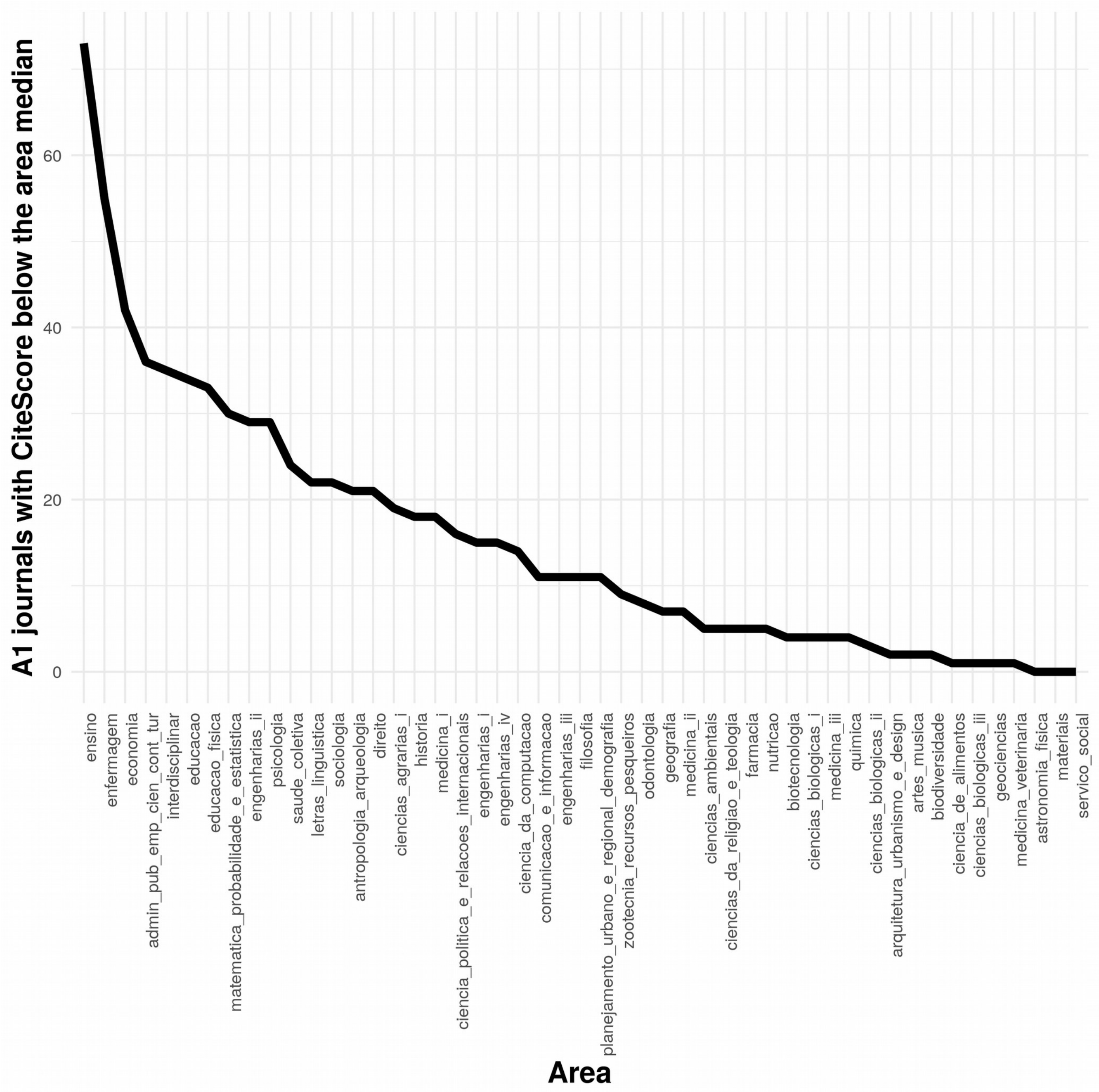


**Figure S4:** Number of journals classified as A1 having a Scopus CiteScore below the median CiteScore for each subject area, across all 49 QUALIS subject areas. Original QUALIS subject area names are shown (as written in their respective classification sheets) but their English translation can be found in Table S1.

**Appendix 1**

The original data containing QUALIS categories and classification criteria for journals in each subject area can be found in the CAPES website:
https://sucupira.capes.gov.br/sucupira/public/consultas/coleta/veiculoPublicacaoQualis/listaConsultaGeralPeriodicos.jsf
To access QUALIS rankings and classification criteria, the user must select a time period and a subject area (highlighted by red arrows). QUALIS rankings and classification criteria are provided as excel and word documents (inside the red box).

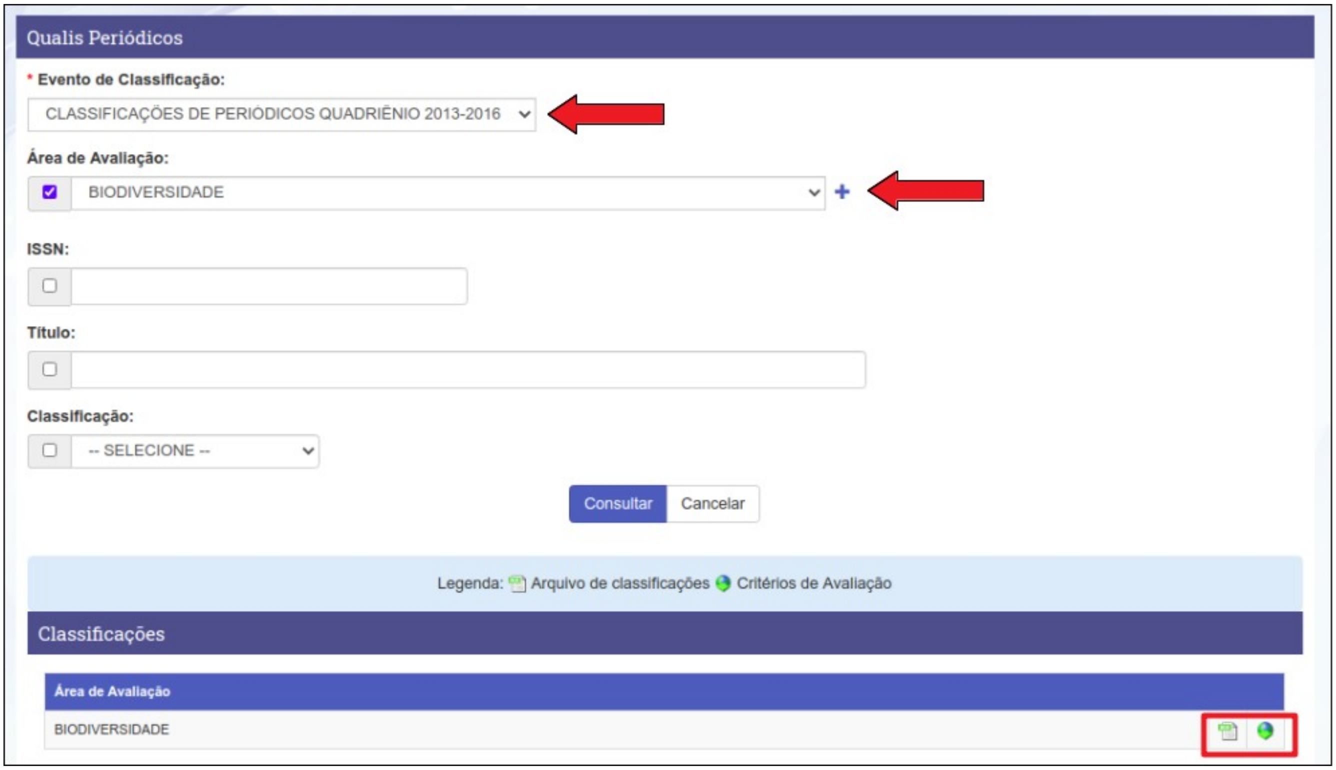
